# The fibrocystin C-terminal domain inhibits Src/STAT3 signal induced cystogenesis of kidney epithelia

**DOI:** 10.1101/2025.08.08.669180

**Authors:** Fatima Hassan, Susanne T. Hahnenstein, Andre Kraus, Bjoern Buchholz, Markus Mukenhirn, Alf Honigmann, Claudia Dafinger, Max C. Liebau, Thomas Pokrant, Jan Faix, Andrea Grund, Dieter Haffner, Wolfgang H. Ziegler

## Abstract

Autosomal recessive polycystic kidney disease (ARPKD) is caused by impaired function of fibrocystin/polyductin (FPC) in collecting duct epithelia resulting in cyst formation. We hypothesized that the membrane-bound C-terminal FPC domain (FPCct) is necessary to suppress cystogensis and facilitate epithelial homeostasis. In ARPKD, cystic kidney epithelia are characterized by a secretory phenotype associated with high intracellular cAMP levels and enhanced STAT3-dependent transcription. Moreover, impaired FPC function may lead to enhanced activation of Src tyrosine kinase, thereby activating STAT3 signaling and its downstream transcriptional activity. To investigate the effects of FPC loss on the cystic epithelial cell phenotype, we used an established principal-like MDCK cell line (pl-MDCK) and studied monolayers in both two and three-dimensional culture. In this *in vitro* model of collecting duct epithelia, FPC-deficient cells showed two-fold elevated basal cAMP levels and enhanced apical secretion leading to three-fold higher luminal pressure. Forskolin-stimulated elevation of cAMP levels triggered enhanced Src-dependent activation of STAT3 resulting in a pronounced cystic phenotype. Notably, expression of wildtype FPCct reduced both STAT3-dependent transcription and the secretory phenotype in knockout epithelial cells. Our data suggest that FPCct interacts with Src kinase at the plasma membrane, thereby reducing Src-mediated STAT3 phosphorylation and limiting STAT3-dependent transcription. Thus, FPCct appears to act like a physiological suppressor of cystogenic signaling, as found in healthy kidney epithelia, that is essential for maintaining epithelial homeostasis. Protein constructs that restore FPC C-terminal function may offer a therapeutic lead to mitigate epithelial dysfunction and slow disease progression in ARPKD.

## INTRODUCTION

Inherited polycystic kidney diseases (PKDs) are disorders caused by biallelic pathogenic variants (or loss) of genes that encode proteins related to the cilia-centrosome complex. The exception to recessive inheritance is the most common disorder, autosomal dominant PKD (ADPKD), wherein disease onset requires a second hit to inactivate the remaining healthy allele. Apart from ADPKD, PKDs are typically rare, monogenetic disorders that involve many different proteins. They belong to the disease family of ciliopathies or ciliopathy-related syndromes, which affect multiple organ systems such as kidneys, liver, and lungs (Hildebrand, 2011; Bergmann, 2018). Despite their relevance and huge efforts, to date there is no curative treatment for inherited PKDs (Liebau, 2022; Zhou, 2023). This is in part due to the lack of appropriate models that allow analysis of disease mechanism, and genetic animal models that at best partially reproduce human disease (Cordido, 2021; Olsen 2019; Ziegler, 2022).

ARPKD is among this group of PKDs and a major cause for kidney failure in infancy and childhood requiring kidney replacement therapy. Complications include bilateral cystic kidneys, lung hypoplasia and ductal plate malformation resulting in congenital liver fibrosis and portal hypertension (Bergmann 2018; Hartung, 2014). The vast majority of ARPKD patients carries biallelic pathogenic variants of the polycystic kidney and hepatic disease gene 1 (*PKHD1*), although variants in other genes can also lead to ARPKD-like phenotypes (Lu, 2017; Bergmann, 2019; Burgmaier, 2021; Halawi, 2023). Pathogenic variants of *PKHD1* lead to functional deficiency of the encoded protein fibrocystin/polyductin (FPC) that localizes to the primary cilia and other cellular membrane compartments. FPC is a large single-pass transmembrane protein of 4074 amino acids that consists of a huge extracellular domain of largely undefined function, a single transmembrane domain and a short cytoplasmic domain, which upon proteolytic release can localize to the nucleus or mitochondria (Kaimori, 2007; Walker, 2023). Proposed functions of FPC protein influence epithelial morphogenesis and homeostasis, with its cell / ciliary signaling affecting transcriptional regulation, mitochondrial function, proteasomal degradation, cell adhesion and proliferation (Bannell, 2024; Harafuji, 2023; Kaimori, 2007; Kaimori, 2017; Walker, 2023; Ziegler, 2020).

While detailed mechanisms of disease onset and progression in ARPKD remain largely elusive, enhanced cell proliferation and cystic expansion of tubular kidney and bile duct epithelia of the liver are proposed to contribute to progressive loss of organ function. It has been assumed that principal cells of the collecting duct promote growth of cysts in PKD (Buchholz, 2011), and that cystogensis arises from defective epithelial homeostasis (Wilson, 1997; Wilson 2004;). Different animal models of ARPKD show enhanced nuclear localization of signal transducer and activator of transcription 3, STAT3, in cyst-lining kidney epithelia (Talbot, 2011; Talbot 2014; Fox, 2024). In addition, the non-receptor tyrosine kinase, c-Src, and STAT3 are consistently activated in many different genetic models of PKD, and both activities are related to enhanced cyclic AMP (cAMP) / protein kinase A (PKA) signaling that is a key feature of cystic kidney epithelia (Zhou, 2022). Thus, targeting (aspects of) cAMP/PKA signaling is central to many pharmacological studies that attempt to develop therapeutic options for inherited PKD (Zhou, 2023). The only FDA-approved drug in the treatment of PKD, specifically its autosomal dominant form (ADPKD), is Tolvaptan, a V2-receptor antagonist that inhibits vasopressin dependent activation of adenylyl cyclase and reduces epithelial cAMP levels. Tolvaptan, however, shows limited effectiveness in reducing cyst progression and is associated with significant adverse side-effects (Torres, 2012).

Here, we used so-called principal-like MDCK (pl-MDCK) cells, a subclone of Madin-Darby canine kidney cells with characteristics of principal-like epithelial cells (Buchholz, 2011; Gekle, 1994), to establish a model of 3-dimensional (3D) epithelial spheroids that mimic tubular kidney epithelia. In this model, single pl-MDCK cells seeded into growth factor-depleted Matrigel divide and, within 3 days, form closed, monolayered epithelial spheroids comprising 24 to 32 cells. The apical membrane of these spheroids facing the lumen is stabilized by an underlying actin belt, a prominent and contractile F-actin structure (Ziegler, 2020; Ziegler, 2022). We used these pl-MDCK spheroids to study the steady-state of water and ion transport across their epithelial barrier and addressed consequences of PKD genetics as well as pharmacological intervention. Lentiviral expression of human wildtype FPCct, a construct of the C-terminal domain, was used to investigate the role of FPC in regulating the epithelial barrier, and to restore FPC function in the *Pkhd1* knockout model of collecting duct epithelia.

## RESULTS

### Principal-like MDCK (pl-MDCK) cells show secretory behavior in presence of elevated cAMP levels

In humans, ARPKD-induced kidney cysts originate from principal cells of the collecting duct and cystic tissue presents with high cAMP levels (Wilson, 2004). To study the effect of enhanced intracellular cAMP levels in our model, wildtype pl-MDCK cells (WT) were seeded into Matrigel in the presence of varying concentrations of forskolin (Fsk) stimulating adenylyl cyclase activity (Figure 1A). pl-MDCK cells formed monolayered spheroids and Fsk (30μM) treatment induced a significant 2.4 fold expansion in lumen size compared to vehicle-treated cells, confirming previous reports in the literature (Buchholz, 2011; Mangoo-Karim, 1989). To detect and quantify enhanced luminal secretion i.e. the cystic phenotype of spheroids, we calculated the “proportional lumen”, which is the ratio of the lumen area to spheroid area at the equatorial plane (Figure 1A) and set a threshold for the proportional lumen of ≥ 0.42 to identify cystic pl-MDCK spheroids. The proportional lumen provides a measure for steady-state water and ion transport into the spheroid lumen and reduces the relevance of the number of cells per spheroid in the analysis of cyst formation. Moreover, a rise in proportional lumen coincides with the appearance of stretched-out epithelial cells indicating enhanced luminal or cyst pressure of spheroids.

**Figure 1.**
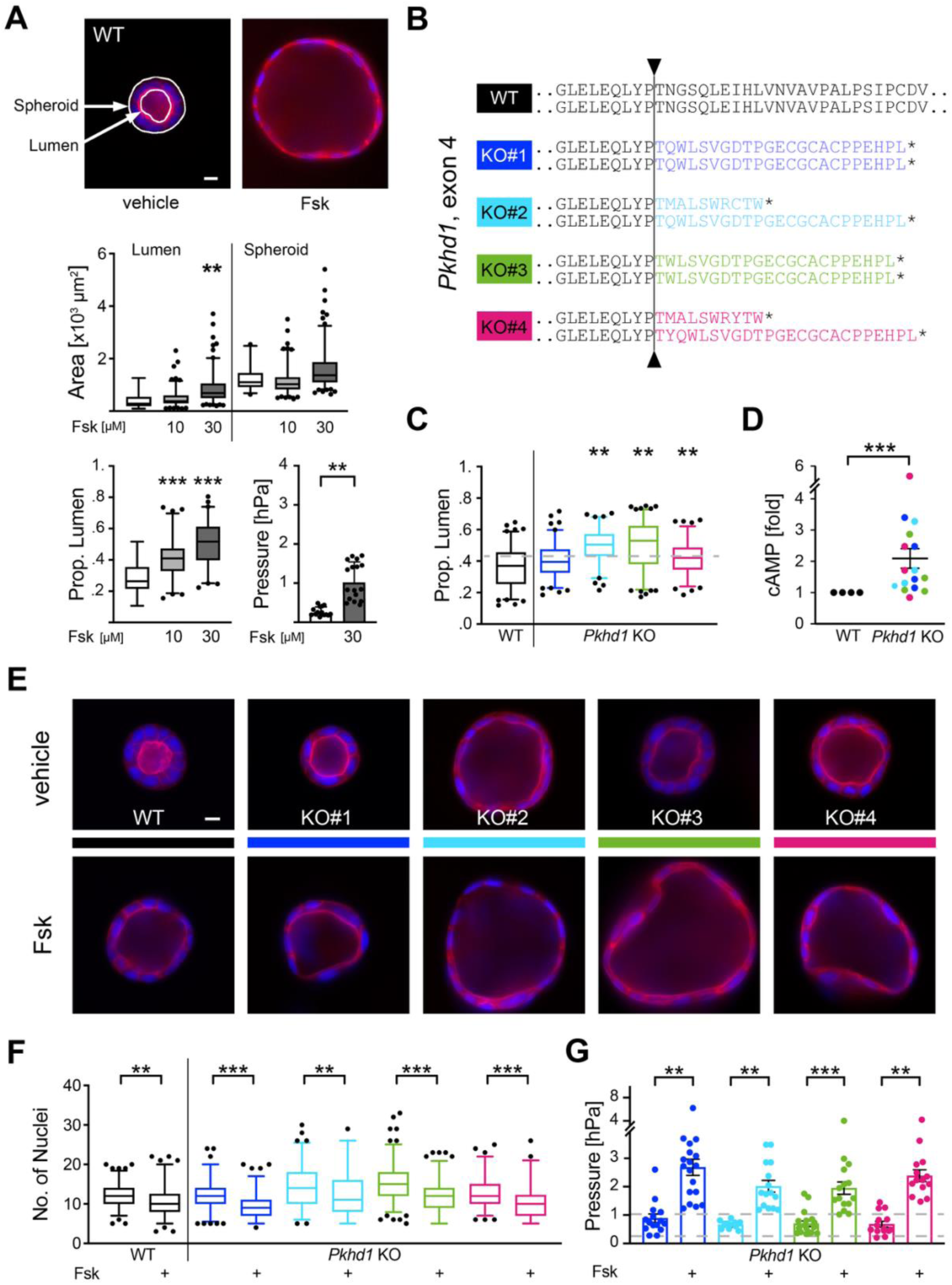
pl-MDCK model for cAMP-induced formation of cystic epithelia. In Matrigel, single pl-MDCK cells form, within 72 hours, monolayered epithelial spheroids an with apical lumen. (**A**) The equatorial plane of spheroids, stained for nuclei (DAPI, blue) and F-actin (phalloidin, red), is used to determine lumen and spheroid area (white arrows, size bar: 10 µm), and to calculate the proportional lumen defined as ratio of lumen area to spheroid area, for each spheroid. Fsk treatment of wildtype pl-MDCK (WT) leads to enhanced luminal secretion, a moderate increase of lumen and spheroid area, and a significant rise of the proportional lumen, as well as hydrostatic luminal (cyst) pressure. (**B**) CRISPR/Cas9-induced frameshift mutations in exons 4 of *Pkhd1* knockout clones, KO#1 to KO#4, leading to loss of FPC protein expression. Note that color code for *Pkhd1* KO clones, as introduced here, is maintained throughout the manuscript. (**C**) Distribution of proportional lumen for pl-MDCK WT and *Pkhd1* KO clones; dashed line: 0.42, and (**D**) cAMP levels, mean ± SEM, in unstimulated epithelia of clones, KO#1 to KO#4, as compared to WT; n=4 independent experiments, non-parametric Wilcoxon. (**E, F**) Fsk stimulation (30 µM) induces cystic epithelial phenotype in spheroids of WT and *Pkhd1* KO clones. The observed rise in proportional lumen (and pressure) results from enhanced ion and water uptake i.e. luminal secretion, and not cell proliferation (see also suppl. Figure S1 B; size bar: 10 µm). (**G**) Hydrostatic luminal pressure observed in individual spheroids (dots), and mean values ± SEM for clones KO#1 to KO#4, vehicle treated or Fsk stimulated; dashed lines (0.24 and 1.02 hPa) indicate mean pressure values of WT spheroids ± Fsk. **Statistics**: spheroid assay (**A, C, F**) >100 spheroids per condition, n=3 independent experiments; box plot with whiskers 0.05/0.95 %, median; non-parametric Kruskal Wallis with Dunn’s post-hoc, **/*** p < 0.01/0.001; pressure (**A, G**) 12 to 18 spheroids per condition, n=3 independent experiments; mixed-effects analysis with Šídàk’s post-hoc; **/*** p < 0.01/0.001.

Upon treatment of wildtype pl-MDCK with Fsk, we observed an increase in the proportional lumen of spheroids by 1.6 and 2.0 fold, with 10 µM and 30 µM Fsk respectively, and a rise in cyst pressure by 4.2 fold in 30 µM Fsk (Figure 1A). Correct apicobasal polarity was established in both Fsk-treated and untreated spheroids, as detected by staining for actin filaments, the apical marker gp135/podocalyxin, and for zona-occludens 1 (ZO-1) to mark tight junctions, thus confirming that epithelial polarity is maintained independent of fluid flow (suppl. Figure S1A). We conclude that pl-MDCK spheroids in Matrigel behave in a manner similar to human kidney epithelia - in terms of their response to Fsk or enhanced cAMP levels, and thus appear suitable as model for kidney epithelia allowing us to study consequences of loss of FPC function in ARPKD.

### Pkhd1 knockout leads to formation of cyst-prone pl-MDCK spheroids

To mimic epithelia of ARPKD patients, we applied the CRISPR/Cas9 method and generated four independent *Pkhd1* knockout (KO) cell clones of pl-MDCK. All clones, KO#1 to KO#4, lack expression of fibrocystin/polyductin (FPC) due to frameshift mutations in exons’ 4 of both *Pkhd1* alleles (Figure 1B). The baseline steady-state behavior of these *Pkhd1* KO clones was assessed determining the proportional lumen of unstimulated spheroids, with values ≥ 0.42 indicating a mild cystic phenotype. When compared to spheroids formed by wildtype cells, *Pkhd1* KO clones KO#2, KO#3 and KO#4 showed a significant increase in their proportional lumen between 10-30%, suggesting enhanced fluid secretion towards the lumen (Figure 1C). Furthermore, enhanced proportional lumen of spheroids was associated with elevated baseline intracellular cAMP levels of clones KO#1 to KO#4 with mean values estimated at 1.7 to 2.3 fold of wildtype pl-MDCK (Figure 1D).

Since a rise in cAMP is a well-established trigger for cyst growth (Yamaguchi, 2000; Wallace; 2011), we recapitulated the cystogenic environment (physiologic environment of PKD cysts) by treating KO spheroids and controls with Fsk. As expected, Fsk stimulation enhanced the proportional lumen by 30-50% in all *Pkhd1* KO spheroids, and also in wildtype spheroids (by 75%), leading to a pronounced cystic phenotype (Figure 1E, suppl. Figure S1B). To also address a possible contribution of cell proliferation to the Fsk-induced cystic behavior, we counted the number of nuclei at the equatorial plane of spheroids. In our experimental setup, Fsk treatment reduced cell numbers by 17-25% in both *Pkhd1* KO as well as wildtype pl-MDCK spheroids, as compared to their vehicle treated counterparts (Figure 1F), therefore, suggesting that Fsk-induced cyst formation is driven by enhanced fluid secretion rather than increased cell proliferation.

Similarly, pressure in *Pkhd1* KO spheroids increased by 2.7 to 3.5 fold upon Fsk stimulation of cAMP levels (Figure 1G, suppl. Figure S2), and mean pressure values observed in spheroids of all clones KO#1-4 were higher as compared to wildtype spheroids, both in vehicle-treated control conditions, by 2.8 to 3.7 fold, and after Fsk stimulation, by 1.9 to 2.6 fold (mixed-effects analysis; comparing values shown in Figure 1A, for WT and in Figure 1G, for KO clones). The epithelial monolayers formed by four independently generated *Pkhd1* KO clones (KO#1 to KO#4) display enhanced cAMP levels, which associate with increased luminal pressure and moderately elevated proportional lumina of *Pkhd1* KO spheroids. Fsk stimulation leads to a strong increase in proportional lumen of spheroids with a luminal pressure about two fold higher in stimulated *Pkhd1* KO as compared to wildtype pl-MDCK spheroids.

### STAT3 activation is enhanced in the Pkhd1 knockout

Elevated phospho-STAT3 (pSTAT3) was reported for ADPKD and ARPKD in human and mouse cyst-lining epithelia and is thought to be involved in cyst growth (Bergmann, 2018; Talbot, 2014; Strubl, 2020). Dafinger and colleagues (Dafinger, 2020) proposed that the short cytoplasmic domain of FPC, the protein encoded by the *Pkhd1* gene, can reduce STAT3 protein activation and nuclear localization by controlling its Src-dependent phosphorylation. Here, we studied pl-MDCK in 2D culture and tested whether STAT3 activation is influenced by the *Pkhd1* knockout. In vehicle-treated control conditions, all *Pkhd1* KO clones showed a slight but consistent increase in STAT3 phosphorylation at tyrosine 705 (pSTAT3-Y705), as compared to wildtype cells, and stimulation with Fsk (10 µM) strongly induced pSTAT3 by 3-4 fold in clones KO#2, KO#3, and KO#4 (Figure 2A, B). Our results support the notion whereby absence of the protein FPC can lead to elevated pSTAT3 levels, and these are further enhanced in cystic tissue by elevated cAMP levels.

**Figure 2.**
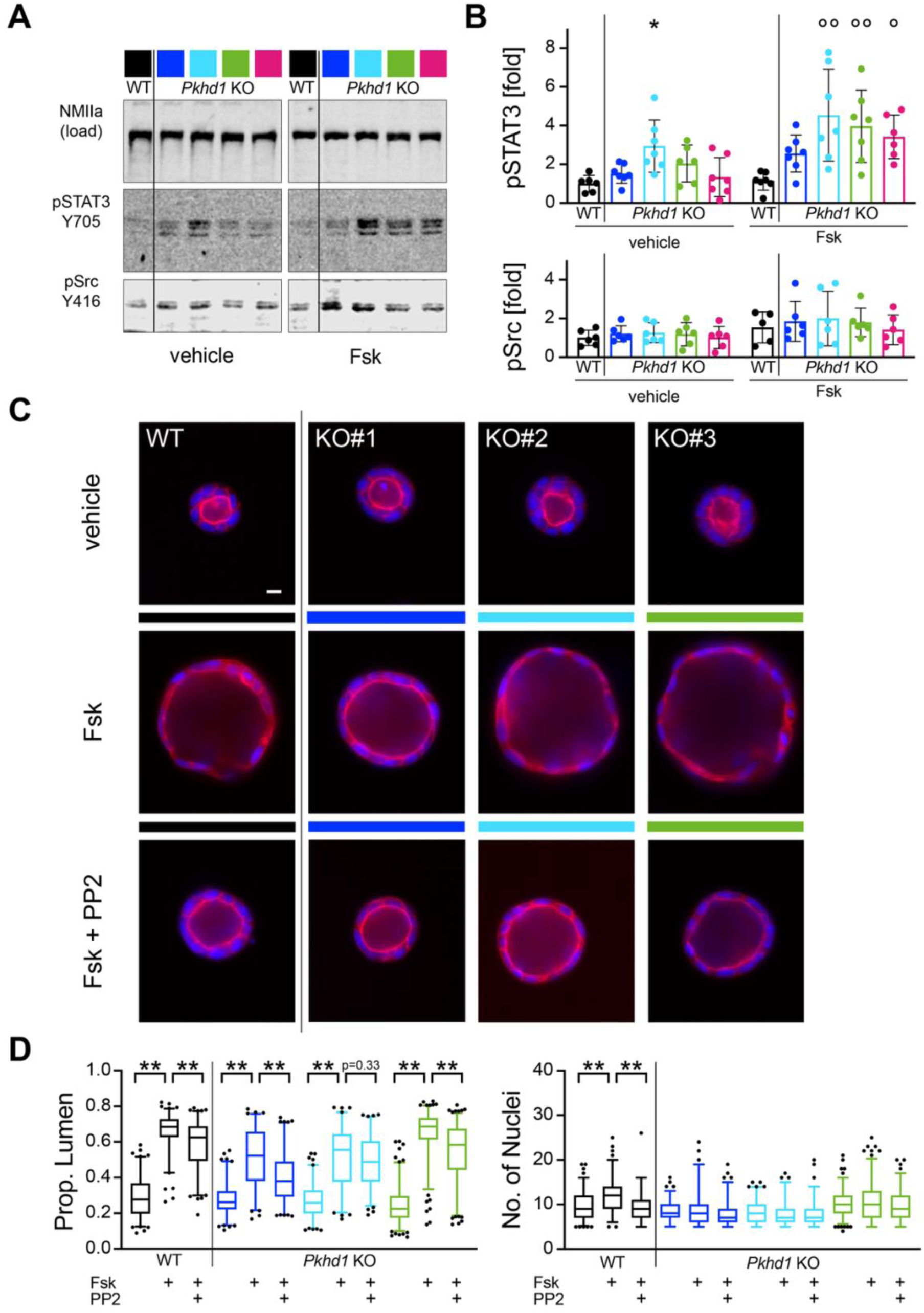
cAMP stimulation leads to enhanced STAT3 phosphorylation and Src-dependent secretory phenotype of *Pkhd1* KO epithelia. Confluent monolayers were analyzed by immunoblot to quantify activating phosphorylation of STAT3 protein at tyrosine 705, pSTAT3-Y705, and activation of Src kinase, pSrc-Y416, in 10 µM Fsk-stimulated pl-MDCK epithelia and controls. (**A**) Representative immunoblot and (**B**) quantitative fluorescence signals of phosphoprotein relative to protein load (non-muscle myosin IIa, NMIIa) for pSTAT3 and pSrc. Impact of Src-inhibition using PP2 (20 µM, 24 h) on Fsk-stimulated cystic pl-MDCK spheroids shown by (**C**) representative images of *Pkhd1* KO and WT spheroids, revealing cell morphology, size bar: 10 µm. (**D**) assessment of proportional lumen (left) and proliferation (number of nuclei in equatorial plane; right) in different treatment conditions, as indicated, confirm reduced luminal secretion of pl-MDCK epithelia, when Src activity is suppressed. **Statistics**: immunoblot (**B**) mean ± SD, n = 3-4 independent experiments; non-parametric Kruskal Wallis with Dunn’s post-hoc; */°,°° p < 0.05, 0.01 relative to vehicle/Fsk-treated WT, respectively; spheroid assay (**D**) >100 spheroids per condition, n = 3 independent experiments; box plot with whiskers 0.05/0.95%, median; non-parametric Kruskal Wallis with Dunn’s post-hoc; ** p < 0.01.

To investigate involvement of the non-receptor tyrosine kinase c-Src, we studied its activation based on phosphorylation of the kinase at Y416 (pSrc-Y416) in both, vehicle-treated and Fsk-stimulated cells. Compared to wildtype pl-MDCK, *Pkhd1* KO clones showed a minor, yet not statistically significant, rise in Src phosphorylation upon Fsk treatment (Figure 2A, B). As Src is involved in multiple signaling pathways, immunoblot analysis of whole cell lysates may not allow detection of specific, spatiotemporally controlled induction of this kinase. Together, lack of FPC expression and cystic conditions (high cAMP) lead to activation of the STAT3 protein, as reported by its enhanced Y705 phosphorylation, and this may result from altered Src activity.

### Src kinase inhibitor PP2 attenuates cyst growth in pl-MDCK spheroids

Several lines of evidence suggested that Src activity contributes to secretory behavior of renal epithelia in PKD (Sweeny, 2008; Talbot, 2014). To establish Src effects on cystic *Pkhd1* KO spheroids, we induced cyst formation by Fsk treatment (72h, 30 µM) and on day 3, suppressed kinase activity by adding a selective Src inhibitor, PP2 (20 µM), for 24 hours, with controls vehicle-treated or induced by Fsk alone. In line with our previous observation, Fsk strongly increased the proportional lumen of both wildtype and knockout cells and subsequent PP2 treatment, i.e. inhibition of Src, counteracted cystic behavior and reduced the proportional lumen of *Pkhd1* KO spheroids, by 12-27%, and to a lower extent also of wildtype spheroids, by 9% (Figure 2C, D).

The estimated average reduction in pressure is 40 Pa or 44% of the Fsk-stimulated increase in luminal pressure (estimation based on the correlation of proportional lumen to pressure; suppl. Fig. S2 C). Furthermore, our data suggest that PP2 reduces cyst growth by limiting luminal secretion in *Pkhd1* KO spheroids and not cell proliferation, as reflected by a rather constant number of nuclei in the equatorial plane of spheroids independent of treatment (Figure 2D). Restriction of Src activity leads to reduced luminal secretion of pl-MDCK epithelia.

### pl-MDCK KO cells express the uncleaved, membrane-bound FPCct protein

Dafinger et al. (Dafinger, 2020) showed that the cytoplasmic domain of FPC can physically interact with Src kinase and thereby inhibit its activation and subsequently STAT3 phosphorylation. To further investigate its impact on STAT3 signaling and cyst formation, we expressed the human wildtype C-terminal fragment of FPC (FPCct), as used by Dafinger and colleagues (Dafinger, 2020), in the pl-MDCK clones KO#1 and KO#3 using lentiviral transduction (Figure 3A). Expression of FPCct comprising a short extracellular domain tagged with V5, the transmembrane (TM) domain, and the cytoplasmic (CP) domain tagged with FLAG, was confirmed based on mRNA detection of V5 and FLAG tags (Figure 3A), and localization of FPCct to the apical plasma membrane (suppl. Figure S3).

**Figure 3.**
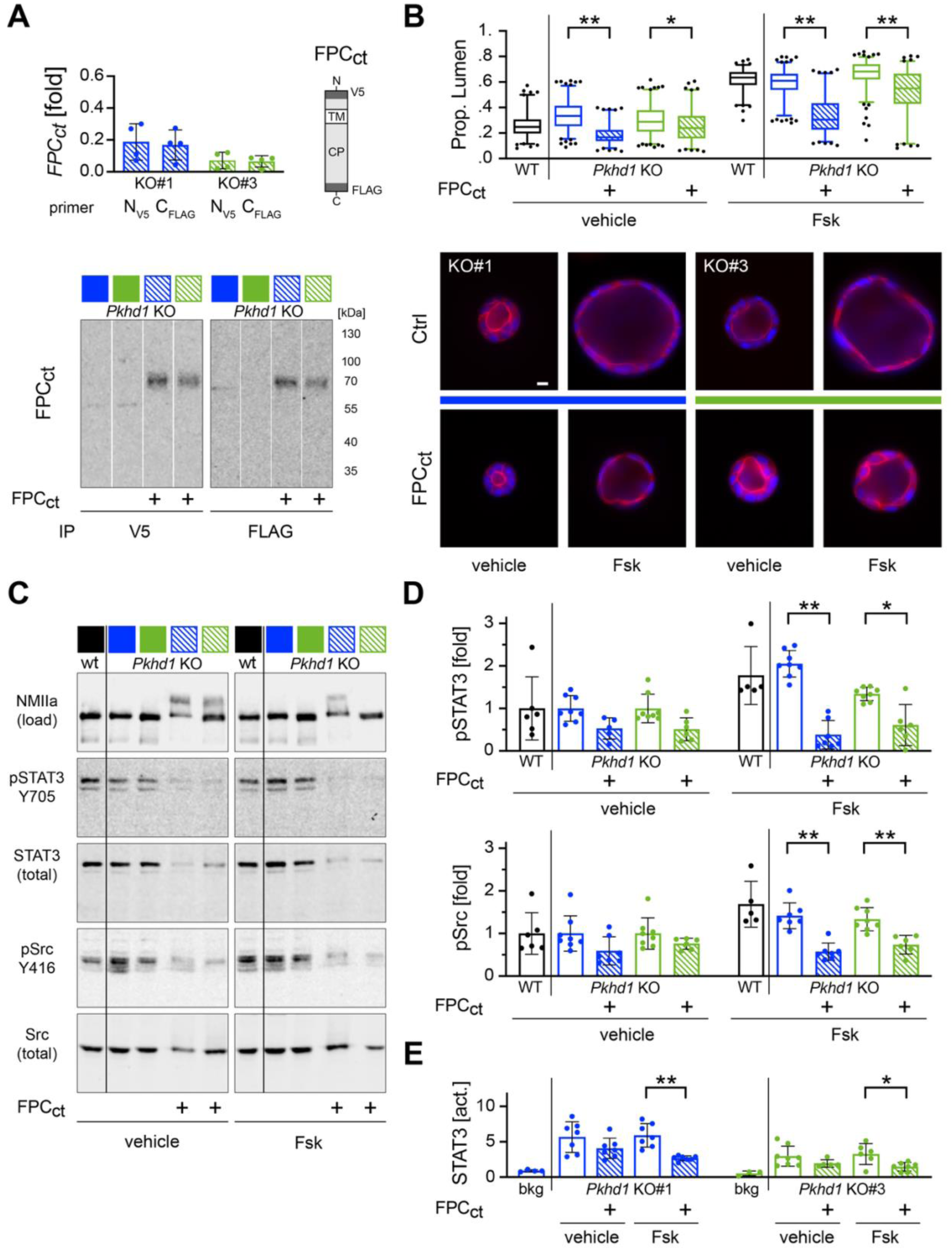
Expression of FPC domain reduces cyst formation, STAT3 activation and STAT3-dependent transcription. (**A**) Transient expression of FPCct, a membrane-bound C-terminal FPC domain construct (cartoon; TM: transmembrane domain; CP: cytoplasmic domain), in *Pkhd1* KO clones, KO#1 and KO#3, as detected by qPCR and immunoblot. Bands of immunoprecipitated (IP) protein reveal full-length FPCct with N- and C-terminal tag, using IP and detection with V5 and FLAG antisera, respectively. (**B**) In spheroids of pl-MDCK clones, KO#1 and KO#3 (upper panel), the proportional lumen is reduced upon expression of FPCct in both vehicle-treated cells and after Fsk stimulation, as indicated, with WT reporting control spheroids; and correspondingly (lower panel), cystic morphology of spheroids is repressed by FPCct expression; size bar: 10 µm. (**C, D**) Expression of FPCct leads to a strong reduction of STAT3 protein and its activating phosphorylation at Y705. In addition, Y416 phosphorylation and activation of Src are suppressed in Fsk-stimulated cells expressing FPCct, as revealed by representative protein bands in immunoblot and quantification of phosphoprotein bands relative to protein load (NMIIa), for STAT3 and Src, which indicates reduced Src/STAT3 signaling. (**E**) Applying the luciferase reporter assay, reduced STAT3-dependent transcription was observed in Fsk-stimulated cells with FPCct expression. Values represent ratio of firefly to renilla luciferase signals, normalized to background (bkg). **Statistics**: spheroid assay (**B**) >200 spheroids per condition, n=3 independent experiments; box plot with whiskers 0.05/0.95 %, median; non-parametric Kruskal Wallis with Dunn’s post-hoc; immunoblot (**D, E**) mean ± SD, n=4 independent experiments; Mann Whitney to compare KO clones ± FPCct expression. */** p < 0.05 / <0.01.

Since full-length FPC was reported to become proteolytically cleaved at the transmembrane domain, releasing a cytoplasmic fragment that translocates to the nucleus (Hiesberger, 2006; Kaimori, 2007), or when further processed, also to mitochondria (Walker, 2023), FPCct protein processing was studied based on immunoprecipitation (IP) with V5 and FLAG antisera. In epithelial monolayers transiently expressing moderate to low protein levels, we observed full-length FPCct protein (70kDa) only, with no evidence of cleaved fragments in the protein pull down (Figure 3A). We conclude that viral expression of FPCct in pl-MDCK monolayers leads to functional protein interactions of the human FPC cytoplasmic protein domain at the apical (or other) membrane(s).

### Expression of FPCct reduces the cystic phenotype of Pkhd1 KO spheroids

Two ARPKD patients with portal hypertension and polycystic kidneys were reported with truncating mutations at the FPC cytoplasmic domain thus revealing its relevance in the disease (Dafinger, 2020). We asked whether expression of FPCct can suppress or even reverse secretory behavior of *Pkhd1* KO epithelia which are deficient for FPC function. To this end, we used lentiviral transduction to transiently express FPCct in KO#1 and KO#3 cell clones and assessed spheroid formation in presence or absence of Fsk stimulation. When expressing FPCct protein, KO#1 and KO#3 spheroids were characterized by reduced lumen and spheroid size and cells showed less stretched morphology, as compared to transduction controls with vehicle or Fsk treatment (Figure 3B).

Similarly, a significant decrease in proportional lumen by 52% and 17%, respectively, was determined for FPCct expressing KO#1 and KO#3 spheroids, already in unstimulated conditions, and by 50% and 19% after Fsk treatment. All differences refer to respective *Pkhd1* KO spheroids with no FPCct expression (Figure 3B). pl-MDCK wildtype spheroids were analyzed in parallel to monitor cyst-prone behavior of *Pkhd1* KO spheroids in the absence of Fsk stimulation and also the effect of Fsk treatment. Our results indicate that expression of FPCct can reduce secretory phenotype of renal tubular epithelia which were induced by (i) genetics, here *Pkhd1* KO or absence of FPC protein, and also by (ii) disease states or interventions leading to high intracellular cAMP levels. The inhibitory effect of FPCct expression observed on the secretion and proportional lumen of Fsk-stimulated *Pkhd1* KO spheroids, corresponds to a calculated mean reduction of 90 Pa (using regression formula; Fig S2C), and thus, was more efficient than inhibition of Src activity using PP2.

### FPCct affects Src kinase and STAT3-dependent activity in Pkhd1 knockout cells

All lentiviral transduced cell clones and controls were used in parallel for spheroid assays and quantitative immunoblots. To further study effects of FPCct expression on Src/STAT3 signaling, epithelial monolayers were extracted and levels of protein expression and tyrosine phosphorylation determined relative to protein load (to control for cell number). Fluorescence-based quantitative immunoblotting showed reduced expression of STAT3 protein, as well as reduced tyrosine phosphorylation of STAT3 and Src kinase at their activity-related sites Y705 and Y416, respectively (Figure 3C, D). Analysis of n=4 independent experiments revealed that FPCct expression in clones KO#1 and KO#3 reduced Fsk-stimulated pY705-STAT3, by 81% and 55%, and likewise pY416-Src, by 60% and 45%, respectively (Figure 3D). Thus, as a consequence of FPCct expression, we observed a considerable reduction of Src and STAT3 activation in confluent epithelia with enhanced cAMP levels.

To directly address STAT3-dependent transcriptional activity, we used a reporter luciferase assay system under control of the STAT3 promoter (Dafinger, 2020). Epithelial monolayers were either vehicle-treated or Fsk-stimulated to enhance Src-dependent STAT3 phosphorylation at Y705 leading to dimerization and nuclear localization. In this setup, expression of FPCct led to a significant decrease in STAT3-dependent reporter activity by 56% and 54%, respectively, in Fsk-stimulated KO#1 and KO#3, and a moderate reduction not reaching significance, in unstimulated cells (Figure 3E). Together, our data suggest a pivotal role of FPCct in the suppression of cyst formation by *Pkhd1* knockout epithelia, which is achieved by limiting the transepithelial ion and fluid flow and by controlling Src-dependent STAT3 signaling, that is a hallmark of PKD.

## DISCUSSION

Our *in vitro* approach studying pl-MDCK epithelia deficient for FPC expression (and function) demonstrates a physiological function of FPC in stabilizing stressed renal epithelia. *Pkhd1* KO epithelia formed by the newly established pl-MDCK *Pkhd1* KO cell clones, are characterized by elevated cAMP levels and increased luminal pressure values of spheroids in Matrigel, showing cyst-prone properties. When cAMP levels in these epithelia are further stimulated by Fsk to mimic a cystogenic environment in the *Pkhd1* KO, expression of FPCct can efficiently reduce Src/STAT3 signaling and mitigate the secretory phenotype of challenged epithelia. The model we propose for FPC involvement in the control of Src/STAT3 signal transduction is summarized in Figure 4.

**Figure 4.**
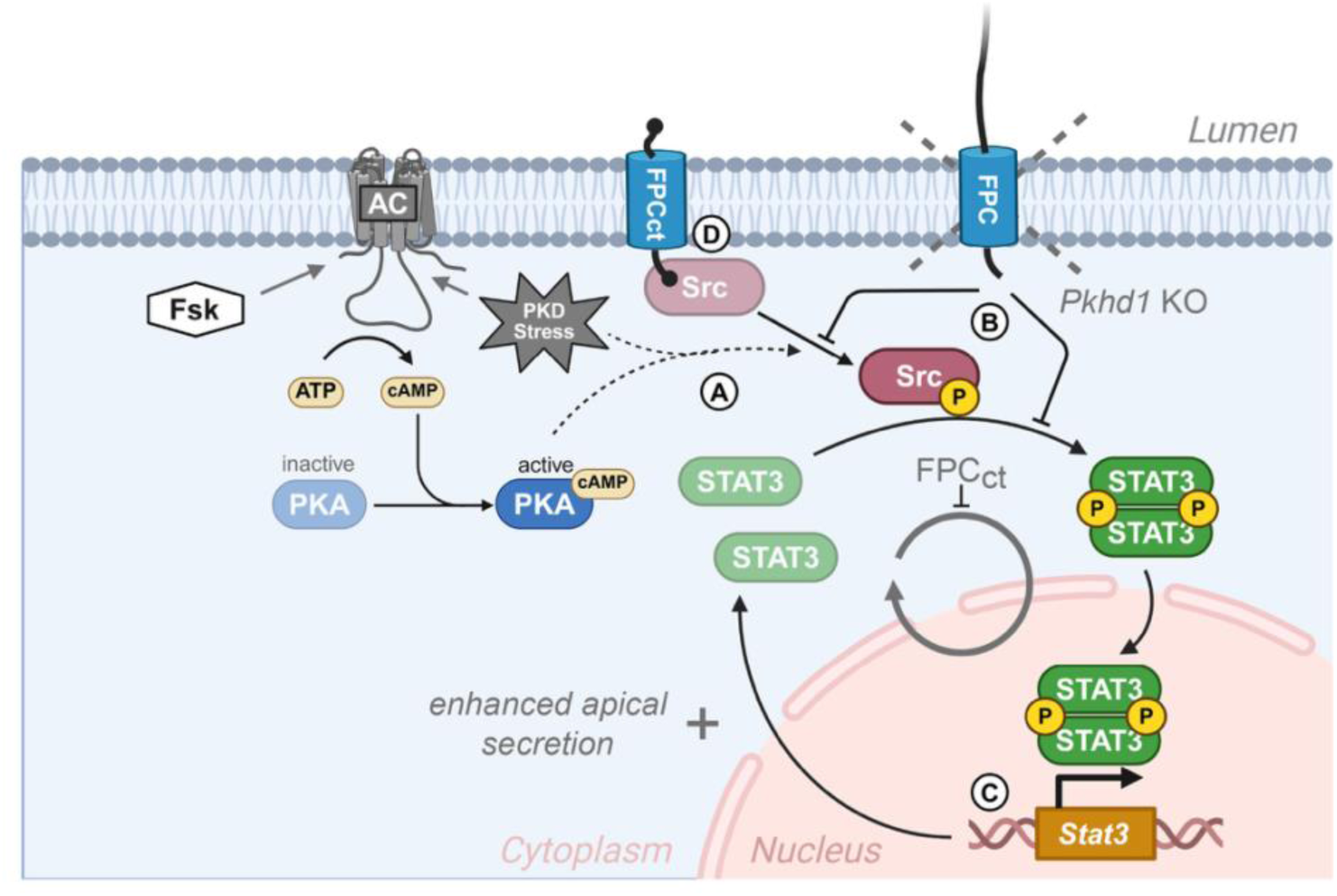
Model of FPC function in stressed epithelia. (**A**) PKD-related epithelial stress induces activation of Src via stimulated cAMP/PKA signaling and other pathways (dotted line). Activated Src phosphorylates STAT3, inducing its dimerization, nuclear localization and STAT3-dependent transcription. (**B**) the FPC cytoplasmic domain interferes with and suppresses both Src activation and signaling, and is absent in the *Pkhd1* KO. (**C**) STAT3-controlled gene expression modifies apical secretion of epithelia and *Stat3* expression. (**D**) Membrane-bound FPCct binds to Src inhibiting its substrate phosphorylation and signaling; for further details refer to Discussion. Created with BioRender.com; **Abbreviations**: AC, adenylyl cyclase; cAMP, cyclic AMP; FPC, fibrocystin/polyductin; FPCct, membrane-bound C-terminal FPC domain construct (with peptide tags at both ends); Fsk, forskolin; P, protein phosphorylation; PKA, protein kinase A; Src, non-receptor tyrosine kinase c-Src; STAT3/*Stat3*, signal transducer and activator of transcription 3 (protein / *gene*)

### Assessment of epithelial function

Principal-like MDCK cells in Matrigel culture constitute a spheroid-based *in vitro* model of collecting duct epithelia that allows study of cAMP/PKA signal-controlled transport of ions and water and cystogenic apical secretion. Expression and control of ion channels, CFTR and TMEM16A, as observed in pl-MDCK, reproduce properties of collecting duct epithelia *in vivo* (Buchholz 2011; Scholz 2022; Zhou, 2022). In this model, the functional readout of epithelial transport activity is achieved using high resolution imaging of closed, monolayered spheroids. The analysis is based on epithelial cell morphology and reproducible quantification of proportional lumen and luminal pressure (Mukenhirn, 2024; Ziegler, 2022). Simulation of cystogenic conditions, applying Fsk stimulation to rise cAMP levels and PKA signaling, leads to a sharp increase of luminal pressure in spheroids by 78 to 180 Pa. This effect falls within the range of membrane pump activity changes reported by Choudhury and colleagues for MDCK epithelia in a microfluidic device (Choudhury, 2022), and represents a 3-4 fold increase in pressure compared to baseline.

Furthermore, changes of spheroid pressure and proportional lumen are strongly correlated (see profile, suppl. Fig. S1), and statistically confirmed changes of mean proportional lumen, as determined in cystic spheroids, with proportional lumen ≥ 0.42, report an increase (or decrease) of luminal cyst pressure by ≥ 40 Pa. These changes are bigger than 24 Pa the average baseline pressure observed for unstimulated wildtype pl-MDCK spheroids. Thus, characteristics of the assay allow sensitive and reproducible assessment of epithelial transport activity in this *in vitro* model. Interestingly, in the experimental conditions of this study, cAMP-induced modulation of proportional lumen and luminal pressure largely occurs independent of detectable stimulation or inhibition of cell proliferation. During the 3-day-period of the assay, we also do not observe any proliferative advantage of the *Pkhd1* KO clones.

### Genetic bias of epithelia

PKD genetics are expected to affect the unstimulated basal transport activity of kidney epithelia (Bergmann, 2018), and consistently, the knockout of polycystin 1 (PC1) led to a reversal of chloride and water transport and apical secretion by MDCK monolayers (Choudhry, 2022). In addition, pl-MDCK cells deficient for PC-1 showed increased cAMP levels, and within a collagen matrix, switched from tubule formation to a cystic epithelial phenotype. Likewise, stimulation of high cAMP levels in wildtype pl-MDCK caused changes in behavior resembling the PC-1 knockout (Scholz, 2022). These observations in genetically modified MDCK cells document that knockout (or functional inactivation) of PKD genes can cell-autonomously affect steady-state properties of kidney epithelia. Similarly, epithelia formed by *Pkhd1* KO clones used in this study, also display cyst-prone characteristics including significantly enhanced cAMP levels, proportional lumen, and luminal pressure, as compared to pl-MDCK wildtype epithelia. These effects occur independent of an external, cystogenic stimulus. When cystic properties are further stimulated by pharmacologically enhanced cAMP / PKA signaling (Zhou, 2022), pl-MDCK epithelia lacking FPC function, KO#1 to KO#4, generate a 2-3 fold higher luminal pressure as compared to wildtype, suggesting enhanced sensitization of ARPKD epithelia towards cystogenic stimuli.

Induction of cAMP / PKA signaling is not necessarily linked to increased epithelial proliferation (Zhou, 2022). While a massive induction of apical secretion is observed in Fsk-stimulated *Pkhd1* KO clones independent of a change in cell proliferation, activating phosphorylation of STAT3, at tyrosine Y705, is upregulated several fold. Enhanced activation of STAT3 was also observed in cyst-lining epithelia of multiple rodent models of PKD (Strubl, 2020), including murine *Pkhd1* knockout (Fox, 2024), as well as, bpk and orpk, genetics that phenocopy ARPKD (Talbot, 2011) and PCK rats (Talbot, 2014). Furthermore, increased activation of c-Src and STAT3 were reported in human cyst-lining epithelia, and elevated pSTAT3 levels were detected in human ARPKD kidney lysates (Dafinger, 2020). Whether increased STAT3 signaling initiates cyst formation or STAT3 merely becomes activated by cystic epithelia is not resolved. The pl-MDCK model of collecting duct epithelia suggests that cystogenic conditions, involving activated cAMP / PKA signaling, lead to enhanced STAT3 activation in the *Pkhd1 KO*, a self-reinforcing consequence of functional FPC protein deficiency, Figure 4.

### Membrane function of FPCct

The relevance of the FPC C-terminal domain function in PKD was challenged by generation of a murine FPC exon 67-deletion model (Δ exon67), which largely lacks the cytoplasmic tail of FPC, yet does not develop any signs of cystic kidney or liver disease (Outeda, 2017). In contrast, two patients with *PKHD1* variants affecting the FPC cytoplasmic tail showed signs of portal hypertension, enlarged cystic kidneys and chronic kidney disease stage G4 in adolescence (Dafinger, 2020). When combined with different PKD genetics, however, the murine FPC Δ exon67 allele enhances cystogensis in the kidney of the PKD1 v/v mice, an ADPKD model with PC1 cleavage defect (Walker, 2023) and mimics effects observed for the FPC knockout in the background of PKD1 variants (Olson, 2019; Walker, 2023). This indicates a critical role of the FPC cytoplasmic domain in the suppression of cyst formation. In these studies, FPC function was attributed to proteolytic processing and release of cytoplasmic peptides that are associated with the nuclear and/or mitochondrial function(s) of FPC (Walker, 2023; Harafuji, 2023). A protein interaction study performed in HEK293T cells reported a further aspect of FPC function. FPCct directly binds to Src and attenuates its activation, which in turn suppresses STAT3 activity (Dafinger, 2020). In pl-MDCK epithelia expressing low to moderate levels of FPCct, we were unable to detect cleavage products of the cytoplasmic domain. Taking into account the membrane localization of the FPCct protein in epithelial monolayers, we suggest that FPCct interacts with Src (and possibly also STAT3) at the apical plasma membrane and thus prevents Src-mediated activation of STAT3-dependent transcription, including expression of the *Stat3* gene itself.

### Targeting Src/STAT3 signaling in PKD

Suppression of Src and STAT3 activities has been proposed and tested in cAMP signal-related PKD models (Zhou, 2022). PKA can directly phosphorylate and activate c-Src. Thus, activation of Src kinase is a convergence point downstream of cAMP/PKA signaling, of several receptor tyrosine kinase pathways (Sweeney, 2008; Zhou, 2022), and also of cell adhesion signaling related to focal adhesion kinase (FAK) (Mitra, 2006). Inhibitor studies confirmed positive effects of suppressed Src activity in different rodent models of ARPKD and reported reduced cyst formation and/or amelioration of PKD and liver cyst disease, for example in PCK rats and bpk mice (Sweeney 2008; Sweeney 2017; Zhou, 2022). A clinical trial for ADPKD based on Tesevatinib, an inhibitor of Src and multiple tyrosine kinase receptors (EGFR, ERBB2 and VEGFR2), proceeded to phase II, but no results have been published. STAT3, the partner of Src, is also phosphorylated and activated by members of Janus family of protein kinases as well as several tyrosine kinase receptors (EGFR, PDGFR and c-MET) (Strubl 2020). Furthermore, cAMP/PKA signaling was reported to increase the effect of cleaved PC1-tail on Src activation of STAT3 (Talbot, 2014). In different PKD models, improved outcome was associated with reduced STAT3 activity (Wang, 2008, Takamura, 2011, Leonhard 2011). So far, there are no clinical trials testing STAT3 inhibition in PKD.

Given the complexity of their regulation, cellular control of Src and STAT3 activities is expected to involve spatiotemporal patterns, and in turn, influences a wide range of cellular properties and functions (Zhou, 2022). A generalized, persistent suppression of one or both protein activities, if possible at all, will cause unwanted or adverse consequences in PKD treatment. Therefore, alternative to directly targeting Src and STAT3 activity, the (partial) reconstitution of FPC function may constitute a suitable strategy to dampen Src/STAT3 signaling and to facilitate or preserve homeostasis of stressed tubular kidney epithelia. This concept of using a protein domain rather than expression of a huge full-length protein, is supported by findings in a murine ADPKD model, where expression of the C-terminal tail of polycystin-1 was sufficient to suppress the cystic phenotype (Onuchic, 2023). Future studies in ARPKD models are required to establish, whether controlled expression of the FPCct protein alone, or in combination with additional therapies targeting cAMP levels, will allow suppression of disease progression and improve epithelial homeostasis.

### Limitations of the study

The study was conducted using an established canine kidney epithelial cell line (pl-MDCK), without access to corresponding kidney tissue of the dog. Furthermore, the study focused on epithelial transport activity and did not account for other factors known to affect epithelial homeostasis, such as tissue fibrosis and inflammatory processes. Failure to detect proteolytically released cytoplasmic fragments of FPCct could be due to their low abundance and/or short lifetime.

## AUTHORS’ CONTRIBUTIONS

WHZ, DH - concept of the study; FH, STH, AK, BB, MM, AH, MCL, JF, AG, DH, WHZ - design and validation of experiments; FH, STH, AK, MM, CD, TP, JF, AG, WHZ - acquisition and analysis of data and samples; FH, STH, AK, BB, MM, AH, CD, MCL, JF, AG, DH, WHZ - interpretation of data and clinical parameters; FH, STH, JF, DH, WHZ - draft and/or revision of the manuscript. All authors read, reviewed and gave their final approval of the work.

## ACKNOWLEDGEMENTS

AH and MM thank the light microscopy facility of the CMCB Dresden for help and support with imaging. STH and FH were supported by the Hannover Biomedical Research School (HBRS). AG, STH, DH, and WHZ received support from the German Network for Early Onset Cystic Kidney Disease (NEOCYST) consortium funded by the German Federal Ministry of Education and Research (BMBF) grant 01GM2203H.

## EXPERIMENTAL MODEL AND SUBJECT DETAILS

Principal-like MDCK (pl-MDCK) cells, a subclone derived from the female cocker spaniel MDCK cell line exhibiting features of principal cells of renal collecting duct, (Gekle, 1994; Buchholz, 2011) were cultured in minimal essential medium (MEM, Sigma-Aldrich, Germany) supplemented with 10 % heat-inactivated fetal bovine serum (FBS, #S1810, Biowest, France), 50 U/ml pencillin, 50 µg/ml streptomycin (Gibco, USA) and 2mM GlutaMax (Gibco, USA). Cells were grown at 37° C, 21 % O_2_ in an incubator with 5 % CO_2._ The cell line was not authenticated.

## METHOD DETAILS

### *Pkhd1* knockout by CRISPR/Cas9

To generate single guide RNA (sgRNA) sequences of 20 nucleotides with high efficiency and minimal off-target effects covering all possible splice variants, gene ID was typed into the CRISPR design tool from Synthego (https://design.synthego.com/#/). The sequence 5’-CACCGAGCAGCTCTATCCTACCAA-3’, targeting exon 4 of the *Pkhd1* gene, was inserted into the BbsI site of plasmid pSpCas9(BB)-2A-puro (PX459) V2.0 (#62988, Addgene, USA) and transfected into pl-MDCK cells (60-70% confluent) using 3 µg plasmid DNA and JetPRIME transfection reagent (#101000027, Polyplus-Transfection S.A, France). After 24 h, the cells were selected with 2.5 µg/ml puromycin (InvivoGen, France) for 4 days and then cultivated for another 24 h in the absence of puromycin. For isolation of clonal knockout cell lines, single cells were subcloned by dilution. Subsequently, the clonal cell lines were expanded and genotyped to verify the *Pkhd1* knockout. To this end, DNA fragments of approximately 700 bp spanning the target site were amplified by PCR from genomic DNA of pl-MDCK control and CRISPR transfected cells using the primers 5′-AAACACAATCCAAGCCCTCCT-3′ and 5′-CTGACCCATCTGTACCAGGG-3′. The resulting amplicons were analyzed by the TIDE sequence trace decomposition web tool (http://shinyapps.datacurators.nl/tide) (Brinkman, 2014). Successful disruption of the *Pkhd1* gene was additionally confirmed by inserting the amplified fragments of respective clones into plasmid pJET 1.2 (#K1231, Thermo Scientific, Lithuania) followed by DNA sequencing (Eurofins, Germany). Four different knockout clones carrying different mutations were selected for the experiments.

### Lentiviral-mediated expression of FPCct

The cDNA sequence of the C-terminal fragment of fibrocystin (FPCct), tagged with V5 at the N-terminus and FLAG at the C-terminus, was subcloned into a lentiviral vector downstream a CMV promoter. Lentiviral vectors were designed, produced and purified by (InSCREENex, Germany). To express FPCct, KO#1 and KO#3 cells (8× 10^5^ cells) were transduced with 9.6 µl lentiviral vector (titer >1× 10^7^ IU/ml) using polybrene transfection reagent (#TR-1003, Sigma, Germany). After 48h, cells were selected for 4 days with 800 μg/ml G418 (Roche, Germany) before being used in the experiments.

### Epithelial cell culture 2D and 3D

#### 2D cell culture

FPCct transduced pl-MDCK cells were seeded into collagen I coated 12-chamber slide (#81201, ibidi, Germany) for 3 days at 37° C, 5 % CO_2_. Cell monolayer was treated with 150 nM MitoTracker Orange CMTMRos (#M7510, Invitrogen, Germany) for 20 min at 37° C, to stain mitochondria, and later fixed with 4 % PFA for 20 min. Cells were permeabilized with 0.25 % Triton-X-100 for 15 min, blocked with 5 % normal goat serum (Merck Millipore, Germany) for 1 h and anti-V5 (1:133; #AB3792, Millipore) or anti-FLAG tag (1:200, #F7425, Sigma) antibody were incubated overnight at 4° C. Cells were later incubated with the secondary antibody goat anti-rabbit IgG (H+L), Alexa Fluor 488 (1:1000; A-11008, Invitrogen, Germany) for 1 h and stained for F-actin with Phalloidin, Alexa Fluor Plus 647 (1:400; #A22287, Invitrogen, US) for 1 h and for the nuclei using DAPI (0.025 µg/ml; Sigma, Germany) for 15 min. Finally, the slide was mounted with Shandon Immu-Mount (Thermo Fisher Scientific, Germany).

#### 3D spheroid assay

*In vitro* 3D assay was adapted from Giles et al. (Giles, 2014) as described (Ziegler, 2022). In brief, single cells (5× 10^3^ for 18-chamber slide or 10× 10^3^ for 8-chamber slide) were resuspended in MEM media containing 2 % FBS and mixed with an equal amount of growth factor reduced Matrigel (#356231; Corning, USA). The mixture was then dispensed either into a µ 8- or 18-well-slide (#80827, #81817; ibidi, Germany), briefly spun at 90x g and allowed to solidify for 30 min. MEM media was added on top of the Matrigel layer and cells were incubated for 3 days at 37° C, 5% CO_2_ to allow spheroid formation. DMSO (0.3 %) (Sigma-Aldrich, Germany) and Fsk (30 µM) (Tocris, UK) were added on day 0 and refreshed after 48h. PP2 (30 µM) (#P0042, Sigma-Aldrich, Germany) was added after 2 days for 24 h. After 3 days of incubation, spheroid structures were fixed with 4 % paraformaldehyde for 30 min (fixing step repeated three times followed by washing with PBS), permeabilized with 0.5 % Triton-X-100 for 20 min and blocked with 5 % normal donkey serum for 1 h. Spheroids were then stained for cell-cell junctions using rabbit E-cadherin (1:500; #3195, Cell Signaling Technology, Germany), apical marker using mouse anti-gp135-Atto550 (1:2000; Antibody Facility (iTUBS, Braunschweig, Germany) and purification/fluorescence-labeling (Hypermol EK, Bielefeld, Germany)). The secondary antibody, F-actin and nucleus stain were used as detailed above in *2D cell culture*. The fluorescence signals were maintained using ibidi mounting media (ibidi, Germany). All experiments were conducted in replicates, with a minimum of three independent repeats.

### Microscopy imaging and spheroid analysis

Images were acquired with Zeiss Axio Observer Z1 microscope, using the 20 x Plan-Apochromat (NA 0.8) objective, the AxioCam MRm Rev.3 camera and the software package AxioVision version 4.8.2 (all from Zeiss, Göttingen, Germany). Filter sets used for imaging were as follow: (1) Alexa Fluor 488—filter set 38 HE, (2) Alexa Fluor 555—filter set 43 HE, (3) Phalloidin-Alexa Fluor 647—filter set 90 HE, and (4) DAPI—filter set 49 (all filter sets from Zeiss).

Spheroid structures were imaged in four colors with 25 z-slice per spheroid with 0.5 µm increment. The structures were analyzed by measuring the spheroid area and the lumen area at the equatorial plane via image J/FIJI, while the number of nuclei was counted manually. At least 100 structures were analyzed per experiment.

### Luciferase reporter assay

To determine STAT3 activity, KO#1 and KO#3 cells (3.5× 10^3^ cells / well) were seeded into a 96-well plate (#24910, Berthold, Germany) and transfected with either: (1) STAT1/3 firefly luciferase reporter plasmid or/and (2) FPCct plasmid, both reported previously (Dafinger, 2020), and (3) Renilla luciferase plasmid pGL4.74 (#E6921, Promega, USA), using JetPRIME transfection reagent (Polyplus-Transfection S.A, France). After 24 h of transfection, cells were treated with Fsk (10 µM) for 1 h and lysed. Firefly luciferase activity was determined using Dual-Luciferase® Reporter assay system (#E1910, Promega, USA) and Tecan Infinite® 200 PRO plate reader (Tecan, Austria). Data were interpreted by dividing firefly luciferase signal over Renilla luciferase signal to normalize for the number of cells and transfection efficiency. Each experimental condition was conducted in triplicate and averaged, with four independent biological repeats.

### cAMP levels

To measure intercellular cAMP levels, wildtype pl-MDCK cells and *Pkhd1* KO clones, KO#1 to KO#4, were grown in a pre-coated assay capture plate (cAMP-Screen Direct System # 4412186; Applied Biosystems, Foster City, CA, USA) and starved in serum-free medium overnight. cAMP measurements were performed using the GloMax-Multi Detection System (Promega, Fitchburg, Madison, USA) according to the manufacturer ‘s protocol. Each clone was seeded in triplicate wells and cAMP levels were averaged. The experiment was independently repeated four times.

### Measurement of luminal pressure in pl-MDCK spheroids

Hydrostatic luminal pressure was measured by quantifying the initial flow rate of fluid exiting the lumen upon laser-induced opening, following the protocol established in a previous study (Mukenhirn, 2024). pl-MDCK wildtype and knockout cell lines were cultured for 3 days, as detailed for the *3D spheroid assay,* to ensure mature cyst formation. To visualize the plasma membrane, cysts were incubated overnight with CellMask™ Orange Plasma membrane stain (5µg/ml) (#C10045, Invitrogen). Lumen drainage was triggered by ablating a single epithelial cell at the cyst midplane using a Zeiss LSM 980 NLO multiphoton system (ZEN blue 3.1), 32X NA 0.8 water objective, equipped with a Spectra Physics (Specta Physics Inc., USA) two-photon laser tuned to 800 nm and 100 % output in the software. A 3 µm-wide line scan across the epithelium, repeated ten times, was sufficient to rupture the cyst membrane and allow fluid escape. The resulting lumen collapse was monitored by acquiring fast time-lapse 3D image stacks at 12-second intervals.

To quantify hydrostatic pressure, lumen volumes were segmented over time using a custom semi-automated pipeline based on LimeSeg in Fiji (https://github.com/Honigmann-Lab-BIOTEC/Tight-junctions-control-lumen-morphology-via-hydrostatic-pressure-and-junctional-tension/tree/main https://doi.org/10.5281/zenodo.12168170). Lumen fluid outflow was modelled as Hagen-Poiseuille flow, describing the movement of a liquid with viscosity η at flow rate Q through a channel of radius R and length L created by the laser cut. The initial flow rate was extracted from the first measurable volume loss.

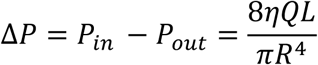

Channel radius and length were previously measured in MDCK-II cysts by imaging the outflow of fluorescent mucin after laser dissection (Mukenhirn, 2024). In the same study, luminal fluid viscosity was determined via fluorescence correlation spectroscopy (FCS) and found to be approximately five times that of water. Since the present study uses pl-MDCK cells - a subclone of Madin-Darby canine kidney - we adopted these previously determined values as representative constants for our system. A total of 12 to 18 spheroids were analyzed across three independent experimental repeats.

### Immunoblot and quantification

To determine pSTAT3 (Y705) and pSrc (Y416) levels, pl-MDCK wildtype or knockout cells were grown until reaching 60-80% confluency and then serum starved overnight. Cells were later stimulated with Fsk (10 µM) or vehicle for 1 h, before being lysed in RIPA buffer containing protease and phosphatase inhibitors (#P8340, #P0044; Sigma-Aldrich, Germany). Protein concentration was determined using BCA assay (#23225, Pierce, Thermo Fisher Scientific, Germany). Proteins (25 µg) were diluted in 1x Laemmli buffer containing 2 mM DTT, heated at 65° C and loaded into 10 % SDS polyacrylamide gel. Protein was transferred onto nitrocellulose membrane, Amersham Protran (#10600003; Sigma-Aldrich, Germany) with Towbin buffer (0.18% methanol, 0.01% SDS) and blocked with 5 % BSA in 50 % TBST and 50 % Intercept (TBS) blocking buffer (#927-60001, LI-COR Biosciences, Germany). Membranes were incubated overnight at 4° C with anti-pSTAT3 (1:500, #9145, Y705, Cell Signaling Technology), anti-pSrc (1:1000, #6943, Y416, Cell Signaling Technology), anti-STAT3 (1:1000, #9139, Cell Signaling Technology), anti-Src (1: 1000, #2110, Cell Signaling Technology), non-muscle myosin IIa (1:1000, #M806, Sigma), anti-V5 (1:1000, #AB3792, Millipore) and anti-FLAG (2.5:1000, #F7425, Sigma). Protein bands were detected using goat anti-rabbit IRDye® 800 CW (1:10,000, #926-32211) or goat anti-rabbit IRDye® 680 RD (1:10,000, #926-68071, LiCOR, USA) via Odyssey Fc Imaging System. Band intensity was measured using image studio software and non-muscle myosin IIa was used as a loading control. Each experiment was repeated at least four independent times, and protein levels were averaged across the replicates.

### Immunoprecipitation

pl-MDCK cells were transduced with FPCct lentiviral construct as detailed above. Cells were lysed on ice in RIPA buffer that contained protease inhibitor and phosphatase inhibitor (P8340, P0044; Sigma-Aldrich, Germany) and centrifuged. Magnetic agarose beads covalently coated with anti-V5 tag (V5-Trap Magnetic Agarose, #v5tma, Proteintech, Germany) and agarose beads covalently coated with anti-FLAG tag (DYKDDDDK Fab-Trap Agarose, #ffa-20, Proteintech, Germany) were diluted and added to the cell lysate, to pull down V5 tag and FLAG tag, respectively. After 1 h of incubation, beads were washed three times and bounded protein was eluted at 95° C in 2x Laemmli, following the supplier protocol (Proteintech, Germany).

### Real-time PCR

Efficiency of FPCct viral transduction was estimated by determining *V5* and *FLAG* mRNA levels in cells via real-time PCR (qRT-PCR). Cells were lysed in RLT (#79216, Qiagen, Germany) buffer containing DTT (Sigma, Germany) and total RNA was isolated using RNeasy Mini Kit (#74106, Qiagen, Germany). RNA (500 ng) was reverse transcribed to cDNA using QuantiTect Reverse Transcription Kit (#205311, Qiagen, Germany) and qRT-PCR was performed using PowerTrack SYBR Green (#A46113, Applied Biosystems, Germany) on QuantStudio™ 6 Pro Real-Time PCR System (#A43161, Applied Biosystems, Germany). Data was analyzed with QuantStudio design and analysis software version 2.5.0. *V5* and *FLAG* mRNA expression were calculated using the formula 2^-Δct^ with *Gapdh* and *Hprt* serving as housekeeping genes. The experiment was conducted in triplicate, with four independent biological repeats.

## QUANTIFICATION AND STATISTICAL ANALYSIS

*Spheroid analysis* data were presented as box plots with 25/75 percentiles, median and whiskers, 0.05/0.95%. Differences among groups were calculated using non-parametric Kruskal Wallis test followed by Dunn’s post-hoc test. *Protein levels, mRNA levels and luciferase activity* were presented as mean ± standard deviation (SD). Differences among groups were calculated using non-parametric Kruskal Wallis test followed by Dunn’s post-hoc test, and between two groups using Mann-Whitney U test. *Luminal pressure and cAMP levels* were plotted as mean ± standard error of the mean (SEM) and analyzed via mixed-effect analysis with Šídák ‘s post-hoc test and non-parametric Wilcoxon test, respectively.

P-value <0.05 was considered statistically significant. Statistical analysis was performed using GraphPad Prism version 9.0; GraphPad Software, San Diego, CA, USA.

