## Supplemental Figures for "The fibrocystin C-terminal domain inhibits Src/STAT3 signal induced cystogenesis of kidney epithelia"

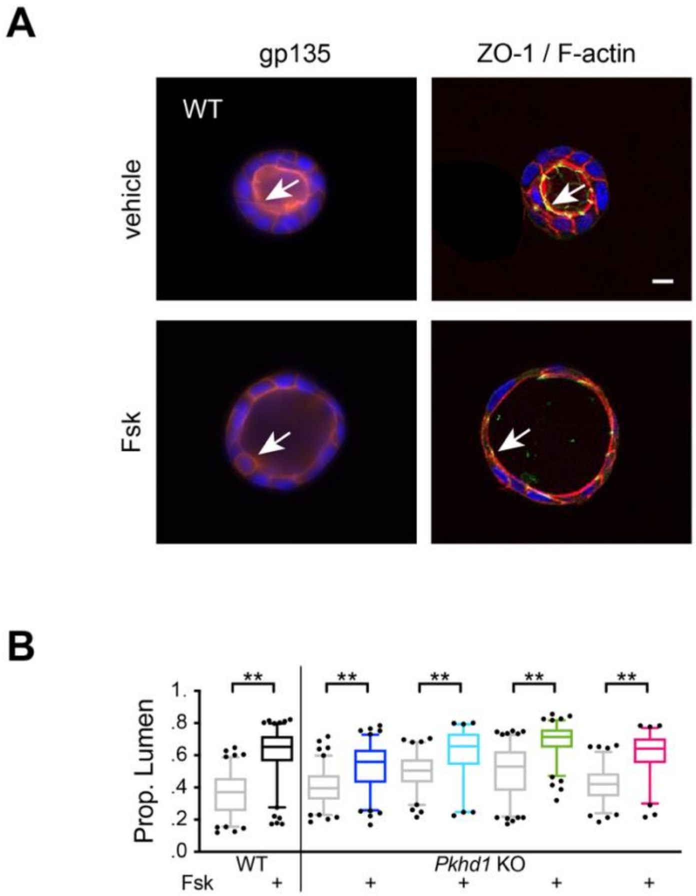


Hassan et al., Suppl. Figure S1

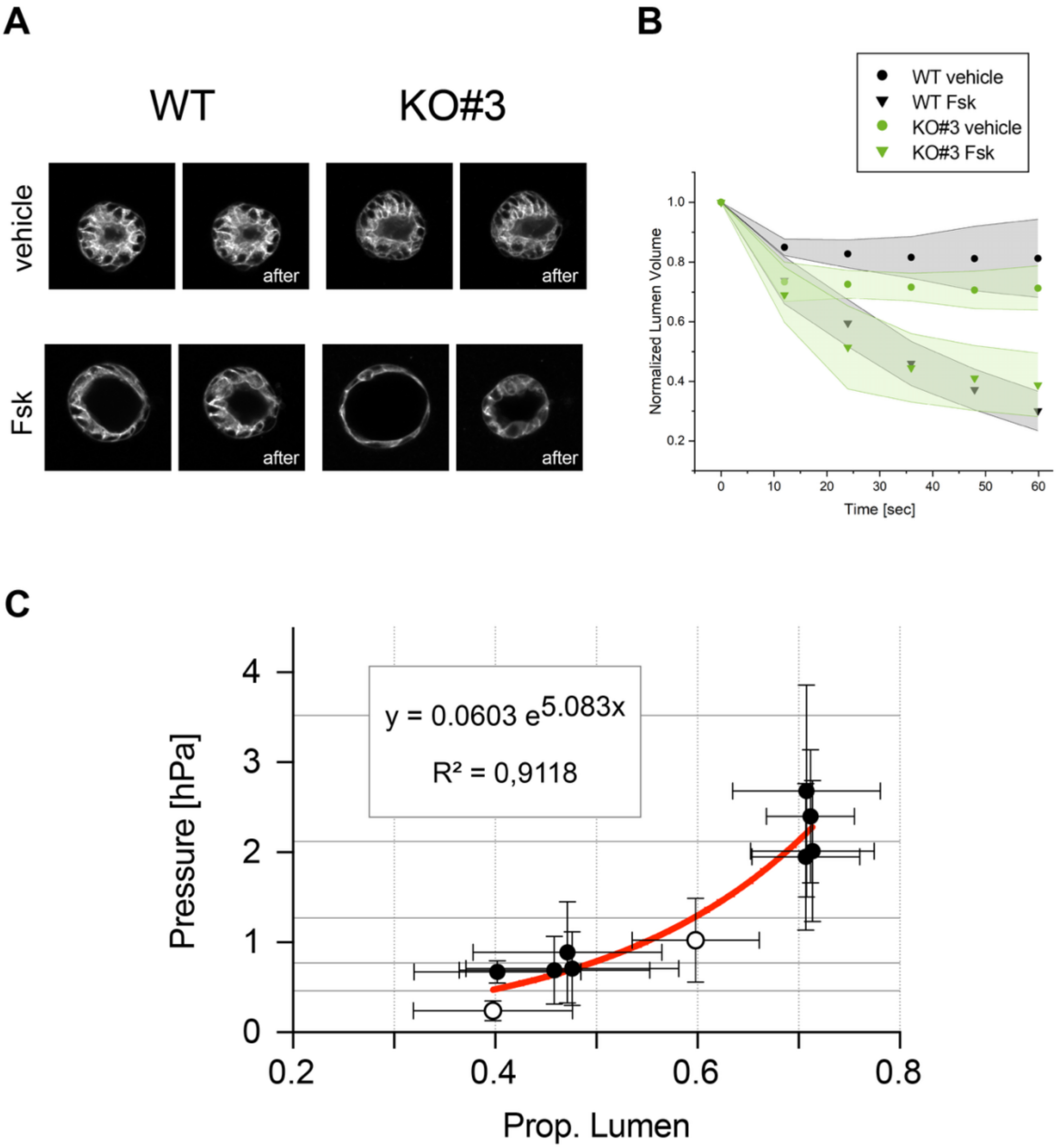


677

678 Hassan et al., Suppl. Figure S2

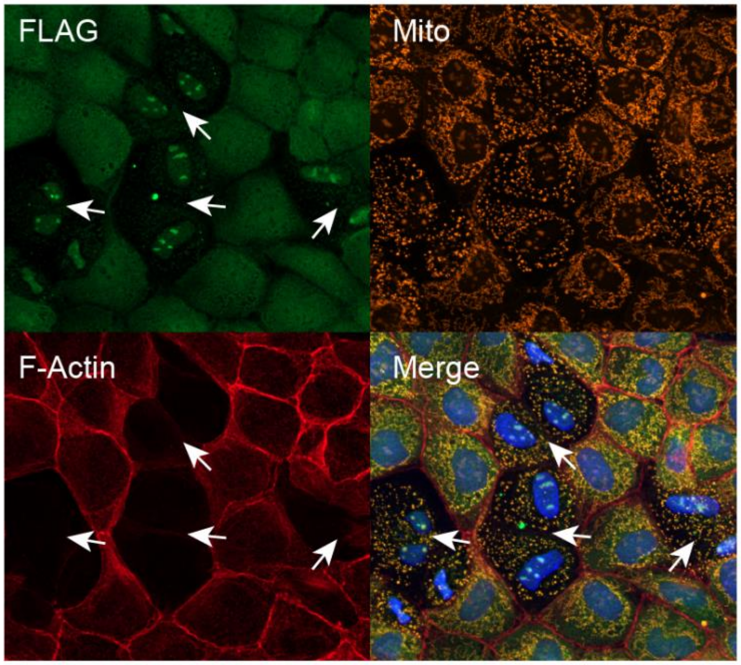


Hassan et al., Suppl. Figure S3

### SUPPLEMENTARY FIGURES

Suppl. Figure S1 **Apical surface and proportional lumen of Fsk stimulated spheroids.** (**A**) Localization of the apical surface (arrows) in pl-MDCK spheroids is not altered by Fsk stimulation (30 µM). Representative images show equatorial plane of WT spheroids stained for apical surface marker podocalyxin / gp135 (red) and nuclei (DAPI, blue), and tight-junction marker, zona occludens (ZO-1, green), F-actin (phalloidin, red) and nuclei (DAPI, blue) as indicated; size bar: 10 µm. (**B**) Proportional lumen of Fsk stimulated WT and *Pkhd1* KO spheroids compared to vehicle treated controls. Controls are shown in grey to indicated that data is used also in Fig. 1C discussing effects of *Pkhd1* KO on unstimulated spheroids; **Statistics**: >100 spheroids per condition, n=3 independent experiments; box plot with whiskers 0.05/0.95%, median; non-parametric Kruskal Wallis with Dunn's post-hoc, ** $p < 0.01$.

Suppl. Figure S2 **Luminal pressure of spheroids and correlation to proportional lumen. (A)** Time-lapse equatorial plane views of pl-MDCK spheroids before and after laser dissection. WT and KO#3 spheroids were either vehicle-treated or stimulated with Fsk (30 µM). Following laser ablation, luminal collapse reflects fluid release, with dynamics dependent on genotype and treatment. (**B**) Exemplary profiles of lumen volume change after laser cutting determined from 3D segmentation of lumina over time. Values are normalized to the initial lumen volume before the cut. (**C**) Correlation of proportional lumen and pressure (hPa=100Pa) using corresponding values of pressure and proportional lumen, means ± SD, of pl-MDCK spheroids, WT (open circles), KO#1-4 (filled circles) with and without Fsk stimulation (n=12-18 spheroids for each circle); equation of exponential regression.

Suppl. Figure S3 **Membrane localization of the FPC domain construct, FPCct** The intracellular distribution of the FPC protein domain is shown in pl-MDCK monolayers based on fluorescent-labelling and detection of FPCct, C-terminal FLAG-tag (green), mitochondria (mito-tracker, orange), F-actin (phalloidin, red) to mark cell-cell junctions, and nuclei (DAPI, blue), individually and in merged image. FPCct localization is characterized by a diffuse staining of the apical surfaces. After cell division (white arrows), when cells are not fully separated (see F-actin), FPC can be detected transiently in perinuclear structures. To document this observation, which is typically rare in densely growing epithelial monolayers, a region of interest with several cell divisions was selected.
